# The sleep EEG envelope: a novel, neuronal firing-based human biomarker

**DOI:** 10.1101/2021.11.16.468788

**Authors:** Péter P. Ujma, Martin Dresler, Péter Simor, Dániel Fabó, István Ulbert, Loránd Erőss, Róbert Bódizs

## Abstract

Sleep EEG reflects instantaneous voltage differences relative to a reference, while its spectrum reflects the degree to which it is comprised of oscillations at various frequencies. In contrast, the envelope of the sleep EEG reflects the instantaneous amplitude of oscillations at specific frequencies, and its spectrum reflects the rhythmicity of the occurrence of these oscillations. The ordinary sleep EEG and its spectrum have been extensively studied and its individual stability and relationship to various demographic characteristics, psychological traits and pathologies is well known. In contrast, the envelope spectrum has not been extensively studied before. In two studies, we explored the generating mechanisms and utility of studying the envelope of the sleep EEG. First, we used human invasive data from cortex-penetrating microelectrodes and subdural grids to demonstrate that the sleep EEG envelope spectrum reflects local neuronal firing. Second, we used a large database of healthy volunteers to demonstrate that the scalp EEG envelope spectrum is highly stable within individuals, especially in NREM sleep, and that it is affected by age and sex. Multivariate models based on a learning algorithm could predict both age (r=0.6) and sex (r=0.5) with considerable accuracy from the EEG envelope spectrum. With age, oscillations characteristically shifted from a 4-5 second rhythm to higher rhythms. The envelope spectrum was not associated with general cognitive ability (IQ). Our results demonstrate that the sleep envelope spectrum is a promising, neuronal firing-based biomarker of various demographic and disease-related phenotypes.

## Introduction

The sleep EEG is a continuous signal reflecting ongoing electrical activity in the brain, and its spectrum reflects the relative contribution of different frequencies to the final waveform. In contrast, the envelope of the sleep EEG estimates the instantaneous amplitude of the signal (typically after filtering for frequencies of interest), and its spectrum estimates the periodicity of all band-limited activity. In other words, the envelope spectrum estimates the typical rhythm at which signal amplitude at certain frequencies waxes and wanes (Figure 1).

**Figure 1.**
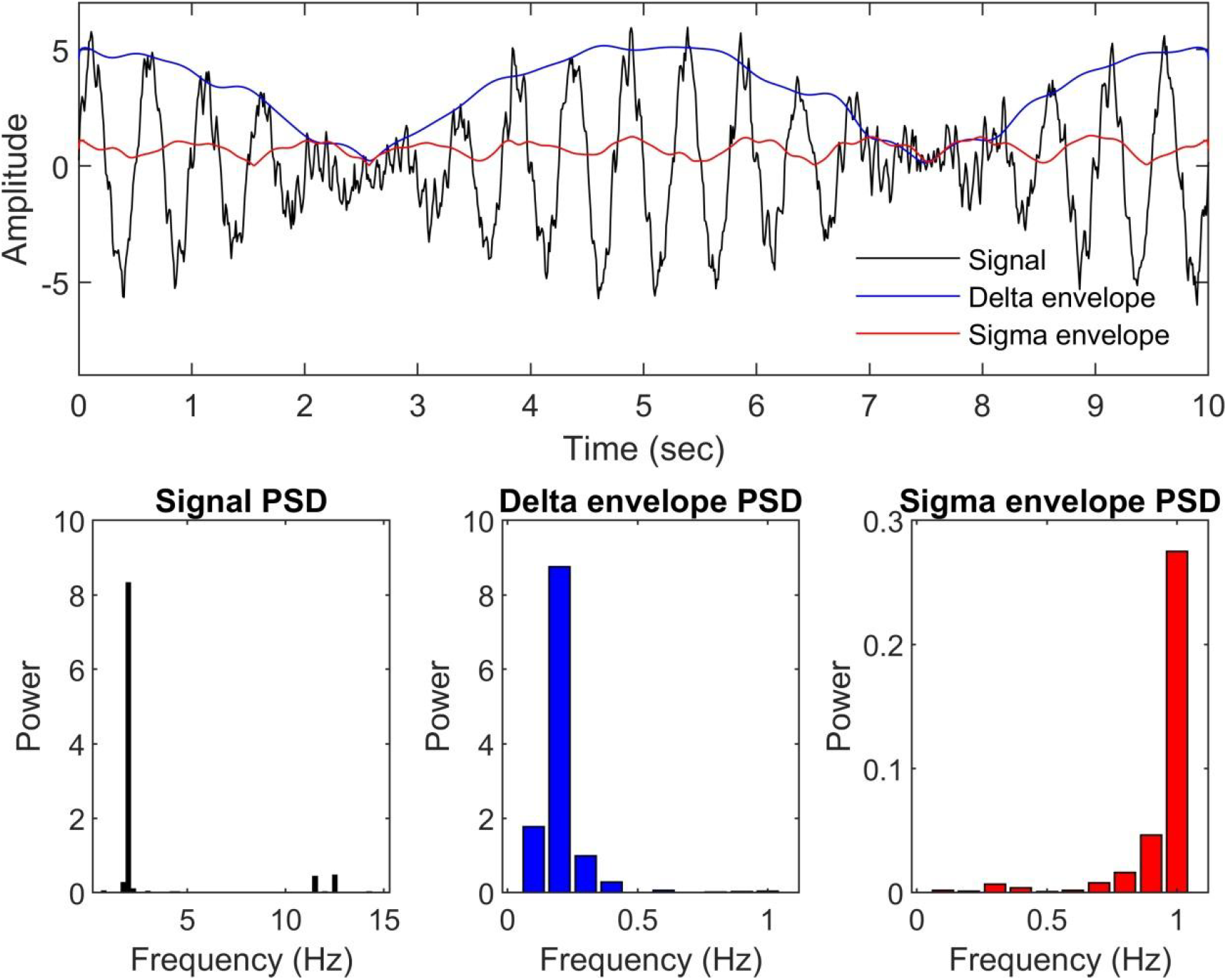
The principle of EEG envelope spectrum analysis. The top plot shows a simulated EEG signal, consisting of the sum of a 2 Hz sinusoid modulated by a 0.2 Hz carrier frequency, a 12 Hz sinusoid modulated by a 1 Hz carrier frequency, and pink noise. Overlain blue and red lines show the instantaneous amplitude or envelope (modulus of the Hilbert transform) of the delta (1-4 Hz) and sigma (10-16 Hz) frequency ranges, respectively. The bottom plots show the power spectral density of the original signal (left) and the delta (middle) and sigma (right) envelopes. Note that the carrier frequencies are accurately recovered from spectral analysis of the envelopes (with some impurities due to added noise and the fact that the modulus of the Hilbert transform of a modulated signal is not fully sinusoidal). The spectrum of the envelope reveals periodic fluctuations in the amplitude of higher-frequency activities.

The spectrum of the sleep EEG signal is one of the best-established general-purpose human biomarkers. First, it shows a fingerprint-like intra-individual stability and inter-individual variability across measurements (Tan et al., 2000; Finelli et al., 2001; Tan et al., 2001; De Gennaro et al., 2005; Reynolds et al., 2018), stabilized mostly by genetic factors (Ambrosius et al., 2008; De Gennaro et al., 2008; Landolt, 2011; Adamczyk et al., 2015). Second, the neural generators of several prominent oscillations contributing to this spectrum have been extensively studied (Steriade, 2003; Csercsa et al., 2010; Nir et al., 2011; Staresina et al., 2015; Gonzalez et al., 2018; Halgren et al., 2019; Fernandez and Lüthi, 2020), often highlighting specific neuronal population assemblies in specific brain structures as their origin. Third, the sleep EEG spectrum has been shown to be a highly reliable marker of age (Sun et al., 2019; Ujma et al., 2019a) and linked to sex (Carrier et al., 2001; Markovic et al., 2020), psychological phenotypes (Steiger and Kimura, 2010; Augustinavicius et al., 2014; Ujma et al., 2017), and a multitude of clinical conditions (Tekell et al., 2005; Fields, 2008; de Haan et al., 2009; Uhlhaas and Singer, 2010; Ozerdem et al., 2011; Kam et al., 2013; Olbrich et al., 2014; Ferrarelli, 2015; Tas et al., 2015; Babiloni et al., 2016; Li et al., 2016; Blinowska et al., 2017; O’Reilly et al., 2017; Purcell et al., 2017). Therefore, the current state of knowledge about the sleep EEG spectrum allows a mechanistic interpretation of how healthy human variability (Tononi and Cirelli, 2014) or disease (Mander et al., 2017) is reflected in neural functioning.

In contrast there has been no systematic research describing the periodicity, generating mechanism or real-life correlates of the EEG envelope, in spite of the fact that as a mathematical function of the oscillations from which the ordinary sleep EEG spectrum is calculated it has the theoretical potential to be an equally promising biomarker with potential incremental validity. On June 1, 2021 we searched PubMed and ScienceDirect with the search terms “eeg envelope spectrum”, “eeg envelope psd” and “eeg envelope power”. Our search returned no relevant papers. We also screened the first 100 Google Scholar hits with these search terms, but also found no relevant papers. Nevertheless, we are aware of some previous studies which do not explicitly assess the spectrum of the EEG envelope but which are still of interest to this field.

An early paper (Kubicki et al., 1986) described that sleep spindles followed each other in periods slightly exceeding four seconds, corresponding to a hypothetical 0.25 Hz envelope oscillation. Two studies (Achermann and Borbély, 1997; Lecci et al., 2017) calculated PSD from short windows, smoothed the resulting power estimates and relied on the spectral analysis of the resulting signal to establish the periodicity of certain frequencies of interest. The first study (Achermann and Borbély, 1997) reported a 20 second (∼0.05 Hz) periodicity for slow waves and a 4 second (∼0.25 Hz) periodicity for sleep spindles, respectively. The second study (Lecci et al., 2017) described a 50 second (∼0.02 Hz) periodicity for both sleep spindles and slow waves, but analyses were restricted to carrier frequencies <∼0.12 Hz. Even slower rhythms have been reported for sleep spindle occurrence (Lázár et al., 2019), replicating the finding of higher amplitude at posterior locations (Lecci et al., 2017). Long-range temporal correlations, especially in the alpha range, were also reported in the wakeful EEG (Linkenkaer-Hansen et al., 2001; Omata et al., 2013). Some studies (Parrino et al., 2000; Terzano et al., 2001) described cyclic alternating patterns (CAPs) as periodic (∼60-90 seconds, 0.011-0.017 Hz rhythms) of both low- and high-frequency activity in NREM sleep, mostly based on visual analysis of the EEG signal.

Several studies of slow EEG rhythms described infraslow oscillations, very slow EEG components typically recorded with direct-current EEG setups which do not impose hardware filter constraints on the lowest detectable frequencies (Vanhatalo et al., 2004; Watson, 2018). Infraslow oscillations are relevant for the study of the EEG envelope because it is a general feature of the sleep EEG that high-frequency rhythms are phase-locked to slower rhythms. As spindles are locked to slow waves and ripples to spindles (Clemens et al., 2007; Clemens et al., 2011; Staresina et al., 2015), virtually all faster rhythms were also shown to be phase-locked to infraslow oscillations (Vanhatalo et al., 2004). Therefore, prominent frequencies in the infraslow oscillation imply prominent frequencies in the envelope of higher rhythms as well.

This preceding literature, however, has not systematically revealed the characteristics of EEG oscillation amplitudes. First, periodicity was estimated only for very specific frequencies, usually slow waves or spindles. Second, somewhat surprisingly, virtually no study used the modulus of the Hilbert transform (or wavelet analogues) as an estimate of instantaneous amplitude, and instead relied on the spectral analysis of smoothed proxies (Achermann and Borbély, 1997; Omata et al., 2013; Lecci et al., 2017; Lázár et al., 2019). Third, as infraslow oscillation studies focused on very slow rhythms, the full range of possible carrier frequencies of interest (especially >0.1 Hz) have not been adequately explored. Fourth, there is little data on the neuron-level generating mechanisms, reliability and real-life correlates of the periodicity of sleep EEG oscillations. In our study, we seek to close this gap by combining data from invasive EEGs of epileptic patients and scalp EEGs of a large sample of healthy participants. We show that the EEG envelope spectrum has at least as many remarkable features as the ordinary EEG spectrum: it reflects neuronal population firing, it is highly reliable within individuals, and it can be used to predict age and sex (but not intelligence) with reasonable accuracy. In line with previous literature, we identify prominent 0.05 Hz and 0.25 Hz rhythms (20 second and 4 second periods, respectively). We also show that the ageing is specifically associated with the loss of 0.25 Hz (4 sec) periodicity of sleep EEG oscillations and the relative amplification of faster rhythms.

## Methods

### Study 1

#### Participants

Sleep electrophysiological data from 13 patients undergoing presurgical electrophysiological evaluation for drug-resistant epilepsy were used. All interventions were approved by the Hungarian Medical Scientific Council and the ethical committee of the National Institute of Clinical Neuroscience. Clinical procedures were not biased for scientific purposes. All patients gave informed consent in line with the Declaration of Helsinki.

#### Electrophysiology

Patients underwent electrophysiological recordings using implanted laminar microelectrodes (IME) and subdural grid and strip electrodes, from which only grids were analyzed (ECoG). Detailed descriptions of these methods are described elsewhere (Ulbert et al., 2001; Csercsa et al., 2010; Ujma et al., 2019b). In brief, IMEs contain 24 serially referenced contacts on a cortex-penetrating pin spaced evenly at 150 µm, capable of detecting extremely local intracortical electrical activity, including neuron population firing, which is represented by high-frequency data (300–5000 Hz) from this source. Multiple-unit activity (MUA), an index of local neuronal population firing, was calculated by rectifying raw data and filtering it with a 20 Hz low-pass filter, according to standard procedure (Csercsa et al., 2010; Ujma et al., 2019b). ECoG was recorded with a sampling frequency/precision of either 2000 Hz/16 bit or 1024 Hz/16 bit depending on the individual patient, and always with a contralateral mastoid reference.

We manually selected seizure-free data with adequate signal quality (indicated by the absence of continuous, broad-frequency artifacts) from all patients. Sleep staging for the selected ECoG data was performed visually on a 20 s basis based on standard criteria (Iber et al., 2007). Since standard scoring criteria are generally only applicable to scalp EEG channels with a full polysomnography setup (including EOG and EMG), we restricted our scoring to the identification of NREM sleep (regardless of stage) and the separation of it from other sleep states and wakefulness, based on the presence of slow waves and spindles. REM sleep, which is difficult to detect using our setup, was not analyzed in Study 1. Artifacts were excluded from ECoG data on a 4 s basis using visual inspection. Only artifact-free data from NREM sleep was considered for further analysis. For analysis, we selected the ECoG channel closest to the IME without epileptiform activity. For the IME, we treated data from poor-quality channels (based on visual inspection) as missing data.

#### Envelope-MUA coupling

In our main analysis in Study 1, we investigated whether fluctuations in the instantaneous amplitude of ECoG oscillations reflected synchronous fluctuations in neuronal population firing within the underlying cortex. For this purpose, we analyzed all artifact-free NREM sleep in each patient, split up into non-overlapping 20-second segments. In each segment, we demeaned ECoG data and used the modulus of the Hilbert transform to estimate the instantaneous amplitude of the following eight frequency bands: low delta (0.5-2 Hz), high delta (2-4 Hz), theta (4-7 Hz), alpha (7-10 Hz), low sigma (10-12.5 Hz), high sigma (12.5-16 Hz), beta (16-30 Hz) and gamma (30-49 Hz). For coupling analysis with each frequency band, we replaced raw MUA data with its moving average calculated from a window of 1/f seconds, where f is the upper limit of each frequency band. The purpose of this transformation was to remove high-frequency components from the MUA signal exceeding the highest frequency at which the corresponding envelope can oscillate.

For each segment and for each frequency band, we estimated the coupling between ECoG envelope and the MUA by calculating 1) the normalized cross-correlation of the two signals implemented with the xcorr() MATLAB function, allowing lags in the [-1 1] second range; 2) the magnitude-squared coherence between the two signals at 0.1 Hz intervals between 0.1 Hz and 1 Hz, implemented with the mscohere() MATLAB function; and 3) the coupling of the amplitude of ECoG envelopes to MUA phases. For this last analysis, we first used the phase angle of the Hilbert transform to estimate the instantaneous phase of the MUA signal. Next, we z-transformed the ECoG envelope signal along the time dimension to standardize amplitude across segments. Finally, we calculated the mean standardized ECoG envelope amplitude (expressed in within-segment SD units) concomitant to MUA data in each of 12 equally spaced phase bins of 30 degrees each. For each patient, we averaged each of the three statistics across all segments to generate a mean value. We used this method to estimate phase-amplitude coupling because traditional methods (Hülsemann et al., 2019), only estimate the preferred phase and overall significance of coupling, whereas we aimed to calculate a more fine-grained estimate. We note, however, that our method is theoretically closest to the Modulation Index (Tort et al., 2008), except we estimate the statistical significance of each histogram bin individually instead of relying on a single, Shannon entropy-based estimate of omnibus significance.

#### Statistical analysis

We estimated the statistical significance of coupling statistics by comparing results to surrogates obtained from random EEG segments. For this, we matched each 20-second ECoG envelope segment with a randomly selected artifact-free NREM MUA segment, calculated cross-correlation, coherence and phase-amplitude coupling and finally an average value across all segments. We performed this analysis 1000 times to generate a null distribution of coupling statistics. An empirical p-value was assigned to each statistic based on actual data, defined as the proportion of surrogate-based statistics more distant from zero.

We calculated unweighted means of all comparable statistics across patients. Similarly, we transformed p-values into standard normal deviates (z-scores) and averaged them across patients, similarly to Fisher’s method of averaging logarithmized p-values (Mosteller and Fisher, 1948). This approach is more conservative and different from ordinary meta-analysis in that it doesn’t add weights to patients based on the amount of data available and it doesn’t increase power over what was originally available in individual patients, so effects which fall short of significance in individual patients do not become significant when data is pooled. Effectively, the alternative hypothesis of this method is that coupling is significantly different from zero in each patient, while in a standard meta-analysis it would be that coupling is significantly different from zero when data from all patients is pooled.

Finally, average standard normal deviates were transformed back to p-values and subjected to correction for false discovery rate using the Benjamini-Hochberg method (Benjamini and Hochberg, 1995) across all lags and IME channels by frequency band (cross-correlation), across frequencies by frequency band and IME channel (coherence), and across phase bins by frequency band and IME channel (phase-amplitude coupling).

### Study 2

#### Participants

We used data from 176 healthy participants (mean age 29.8 years, SD 10.66 years, range 17-69 years; 95 males) from a multi-center database of the Max Planck Institute of Psychiatry (Munich, Germany) and the Psychophysiology and Chronobiology Research Group of Semmelweis University (Budapest, Hungary) (Ujma et al., 2014; Ujma et al., 2019a) was used in this retrospective study. We used participants with available cognitive test scores (Raven’s Advanced Progressive Matrices, the Culture Fair Test and/or the Zahlenverbindungstest [a trail making Test]). Test scores were always expressed as IQ scores with a population mean of 100 and a standard deviation of 15, and if multiple tests were available from a single participant, the scores were averaged (see the first publication of the dataset (Ujma et al., 2014) for details).

Study procedures were approved by the ethical boards of Semmelweis University, the Medical Faculty of the Ludwig Maximilian University or the Budapest University of Technology and Economics. All participants were volunteers who gave informed consent in line with the Declaration of Helsinki. According to semi-structured interviews with experienced psychiatrists or psychologists, all participants were healthy, had no history of neurologic or psychiatric disease, and were free of any current drug effects, excluding contraceptives in females. Consumption of small habitual doses of caffeine (maximum two cups of coffee until noon), but no alcohol, was allowed. Six male and two female participants were light-to-moderate smokers (self-reported), and the rest of the participants were non-smokers. Further details about participant selection criteria and study protocols can be found in the studies reference above.

#### Polysomnography

All participants underwent all-night polysomnography recordings for two consecutive nights, and data from the second night was used for all analyses. Scalp EEG electrodes were applied according to the 10-20 system and referenced to the mathematically linked earlobes. Impedances were kept at <8kΩ. EEG was sampled at 250 Hz for 115 participants, 249 Hz for 29 participants and 1024 Hz for 15 participants, always resampled at 250 Hz. Sleep EEG was visually scored on a 20 second basis according to standard criteria (Iber et al., 2007). A visual scoring of artifacts was also performed on a 4 second basis. EEG preprocessing was implemented in Fercio’s EEG (©Ferenc Gombos, Budapest, Hungary). Further details about the technical details of the sample can be found in the first publication of this dataset (Ujma et al., 2014).

#### Envelope spectra

We used two-way least-squares filtering (implemented in the MATLAB EEGLab function eegfilt()) to filter the sleep EEG of each channel of each participant to the following eight frequency bands: low delta (0.5-2 Hz), high delta (2-4 Hz), theta (4-7 Hz), alpha (7-10 Hz), low sigma (10-12.5 Hz), high sigma (12.5-16 Hz), beta (16-30 Hz), gamma (30-49 Hz). The envelope of each of these frequency bands was calculated using the modulus of the Hilbert transform, resulting in eight signals per participant and channel. We used discrete Fourier transform (DFT, implemented in the MATLAB EEGLab function periodogram()) to estimate the power spectral density (PSD) of the envelope using rolling, overlapping 100 second windows (with 20 second steps and thus an 80 second overlap). The envelope signal in each window was demeaned, detrended and Hamming-windowed before DFT. PSD was estimated between 0.01 Hz and 4 Hz with 0.01 Hz increments for each sampling window, and an average PSD across windows was calculated for each participant, channel and frequency band.

Because fluctuations in the envelope of the EEG signal are expected to take place on a much longer timescale than fluctuations in the signal itself, very low frequencies of the envelope spectrum are of particular interest, but their estimation is only possible with sampling windows much longer than those used to estimate the ordinary power spectrum. This introduces a particular problem when dealing with artifacts. In case of the ordinary power-spectrum, which is estimated using many sampling windows each only a few seconds long, the loss of a few sampling windows due to the presence of artifacts only results in the loss of a comparatively small fraction of the total signal. In case of the envelope spectrum, however, totally discarding a 100-second sampling window due to a presence of a relatively short artifact may result in an unacceptable amount of signal loss. Therefore we used a colliding window method (Figure 2, Panel A) to deal with artifacts. When the 100-second windows sampling the signal in 20 second steps encountered a segment marked as an artifact, they were progressively shortened to end before the artifact, until a minimum sampling window length of 20 seconds was reached. At this point, the sampling window skipped the artifact segment and re-started at its original 100-second duration afterwards. PSD from the shortened windows was calculated and used as usual, but PSD estimates of the frequencies below 1/L Hz were discarded and in the calculation of the average PSD data from this window was under-weighted by 1*L/100 (L in both cases refers to the length of the window in seconds). In order to avoid over-sampling of data before artifacts, all envelope signals were sampled both in the forward and backward direction, starting the 100 second windows from the beginning and the end of recordings, respectively. The colliding method ensured a minimum signal loss of 20 seconds instead of 100 seconds in case of artifacts.

**Figure 2.**
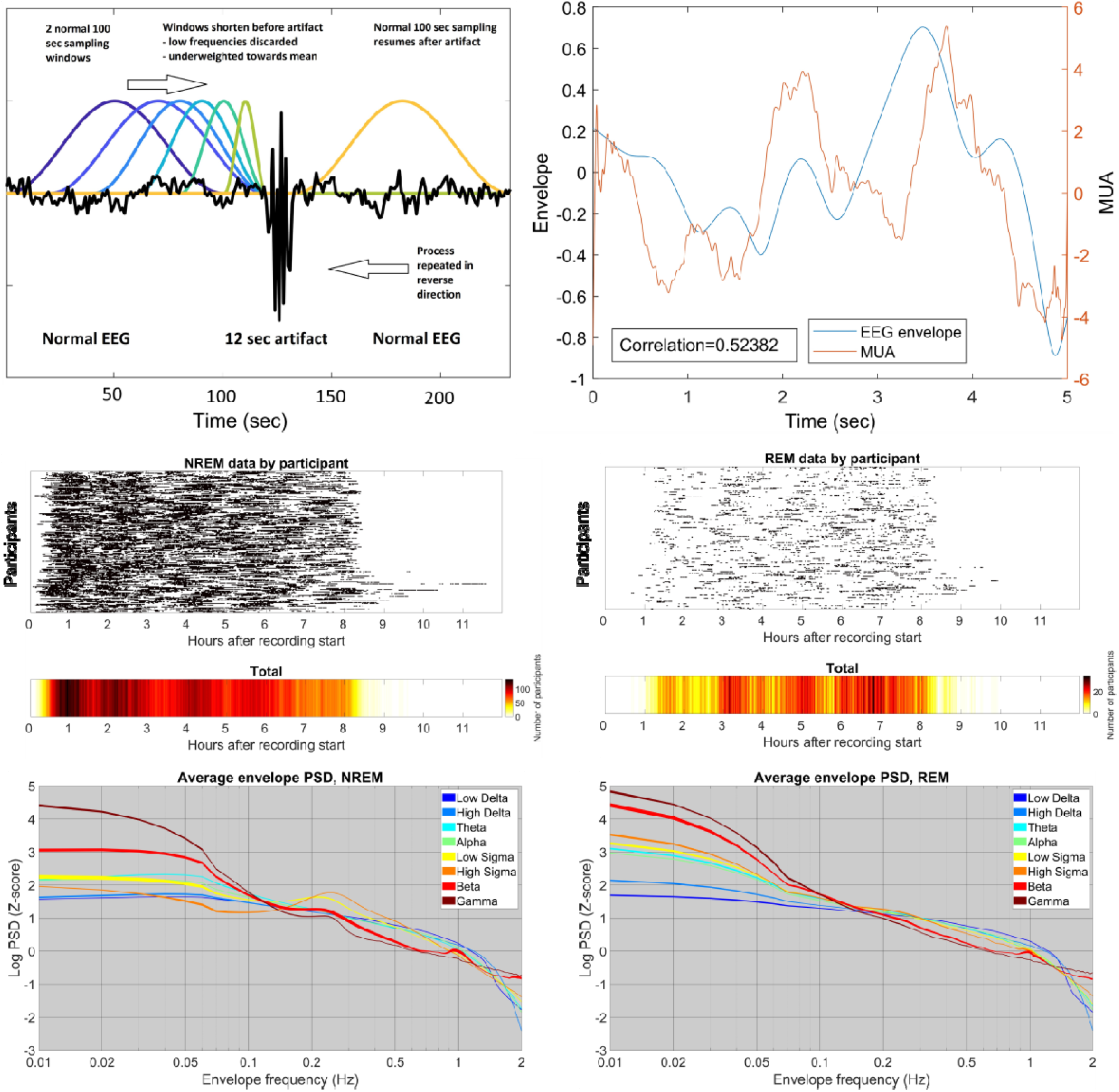
An illustration of EEG envelopes, the colliding window method and its results. **Panel A** illustrates the colliding window method. **Panel B** shows a single epoch of illustrative envelope and MUA data (ECoG low delta envelope and smoothed MUA from the fifth IME channel located in cortical layer III). The Pearson correlation of the two signals is shown for reference. Both the ECoG envelope and the MUA is detrended and demeaned, but not z-transformed. **Panel C** shows the distribution of available sleep data after artifact rejection using the colliding window method. For each participant, black lines mark the data segments used in analysis. The lower panel shows the total number of participants with available data as a function of time after recording start. **Panel D** illustrates the log-transformed envelope spectra. All data was z-transformed by frequency band to eliminate mean differences. The frequency axis is shown on a log scale to enhance the low frequency ranges which are of particular interest. Note spectral peaks at ∼0.05-0.06 Hz, ∼0.25 Hz and ∼1Hz, the latter most prominent in the beta range.

The resulting average envelope PSDs were smoothed using the Savitzky-Golay method with a 10-degree polynomial, 10-base log-transformed to normalize variances for linear statistics and z-transformed across frequencies within participant and channel to eliminate the effect of between-participant differences in raw EEG signal voltage. Participants with abnormal PSDs (based on visual inspection) on any channel in any frequency band were removed from analyses concerning that frequency band (N=1-3 participants per frequency band).

Envelope frequencies up to 2 Hz (that is, fluctuations in EEG amplitude with up to two cycles per second) were considered for analysis. Figure 2 illustrates the colliding window process, the amount and temporal position within the night of available artifact-free data and the average spectra. Detailed individual spectra are available in the Supplementary Data.

#### Statistical analysis

Even-odd reliability was computed by calculating the average PSD for each individual twice, using even and odd numbered sampling windows separately. Because sampling windows were up to 100 second long and overlapped by 20 second steps, only every fifth sampling window was used to avoid non-independent data. The reliability of the PSD in each frequency band, on each channel and at each frequency was estimated by the intraclass correlation coefficient (implemented as Pearson’s correlation coefficient with pooled standard deviations) between the two measurements. For split-half reliability, we also calculated the average PSD for each individual twice, using the first and last 50% of all available sampling windows separately. Because the intraclass correlation coefficient is sensitive to mean differences and we expected mean signal voltage to systematically change between the first and second halves of the night, we computed split-half reliability using the ordinary Pearson correlation instead. Although reliability is generally defined as the square root of the correlation between repeated measurements because they are both expected to be equally affected by unreliability (Schmidt and Hunter, 2014), we used the more conservative and more easily interpretable unsquared coefficients.

For multivariate predictions, we used elastic net regression implemented in the MATLAB lasso() function. Elastic net regression is an iterative learning algorithm which seeks to maximize the predictive value of a large number of potentially correlated predictors by introducing a penalty term for complexity. Elastic net regression is able to fit reliable models in samples where OLS regression would be underdetermined given the large number of predictors and the small sample size. Technical descriptions (Tibshirani, 1996; Zou and Hastie, 2005) and practical implementations (Krapohl et al., 2017; Lello et al., 2017), including in sleep EEG analysis (Ujma et al., 2019a) are available in the literature. We used 5-fold cross-validation and an L1-L2 regularization mixture set at alpha=0.5 for elastic net regression models. All envelope spectral values between 0.01–1 Hz from all spectral bands (800 variables in total) were used as predictors and age, sex (here treated as a continuous variable (Gomila, 2021)) and IQ were used as dependent variables. These models were fitted independently using data from each electrode (18*3=54 models in total). One eighth (N=22) of the sample was retained as a validation sample, and the models were trained on the remaining participants (N=154, including the cross-validation samples used to ensure robust regression coefficients). The models resulting from training were used in the fully independent validation sample to check performance. The validation sample was selected by ordering participants by the values of the dependent variable and taking every 8^th^ individual to ensure maximal variance.

All analyses in all studies were implemented in MATLAB 2018a.

## Results

### The envelope reflects cortical neuronal firing

In Study 1, we correlated the envelope of EEG signals measured at the cortical surface with neuronal firing patterns measured from within the adjacent cortex. We found that in all frequency bands and across the entire cortical mantle, the envelope of the surface signal reflected firing patterns, with a typical average magnitude-squared coherence value of 0.15–0.2. The pattern of coupling was different as a function of frequency range (Figure 3).

**Figure 3.**
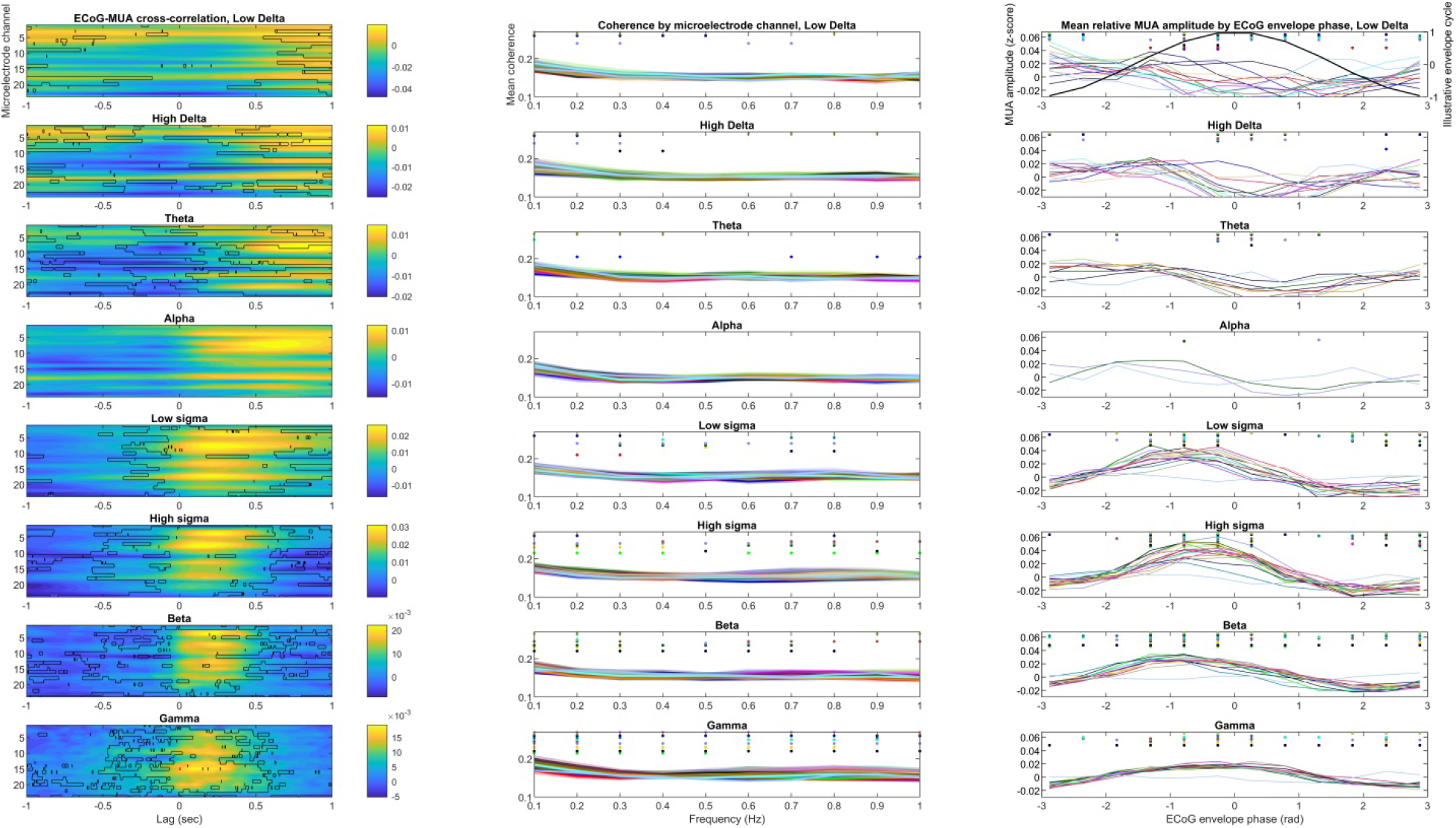
Coupling between EEG envelope in the cortical surface and MUA within the adjacent cortex in NREM sleep. **Panel A:** cross-correlation of the two signals. The horizontal axis indicates time lags, the vertical axis indicates IME channel (N=23, deeper channels shown at the bottom), while the color axis indicates correlation coefficients. Black outlines show statistically significant results after FDR correction. **Panel B:** magnitude-squared coherence between the two signals. Overlain lines represent individual IME channels. Because of the large number of channels and no substantial between channel differences, no particular pattern in color coding was used. Dots indicate statistical significance after FDR correction on the corresponding channel. Deeper channels are shown at the top. **Panel C:** Mean MUA amplitude (in within-segment z-scores) by ECoG envelope phase bins. Dots indicate statistical significance after FDR correction on the corresponding channel. A sinusoid is overlain in the low delta subplot for illustration. On panel B and C, for better visibility only IME channels are shown where at least one data point reached significance.

In the delta through the theta range, MUA was lowest during or slightly after the peak of the envelope, also reflected by the fact that the largest cross correlations were negative and observed when a zero to slightly negative MUA delay was added. In the sigma through gamma ranges, MUA was highest during the ascending phases of the envelope, also reflected by the fact that the largest cross-correlations were positive in case of a positive MUA delay. The alpha range exhibited an intermediate pattern with only a few IME channels reaching significance.

Findings in individual patients are available in the Supplementary Data. We note that in 3 patients the envelope-MUA coupling was absent or restricted to very specific channels. As data issues (problems with synchronization in case of an absent coupling, and poor MUA data quality in case of both absent or spatially restricted coupling) is a possible explanation for this pattern, we re-analyzed coupling excluding these three patients. Results were virtually identical even in this case (Supplementary figure S1)

### The envelope spectrum is stable within individuals

The ordinary sleep EEG spectrum is known to have a trait-like quality by being stable within individuals, but vary between individuals (Finelli et al., 2001; De Gennaro et al., 2005). In the absence of multiple recordings from participants, in Study 2 we assessed the trait-like nature of the envelope spectrum by calculating even-odd reliabilities (intraclass correlation coefficients between the spectral densities calculated separately from the even or odd numbered sampling windows of the same individual) and split-half reliabilities (Pearson correlations between the spectral densities calculated separately from the first and last 50% sampling windows of the same individual).

Based on this analysis, the envelope spectrum was highly trait-like in NREM sleep and moderately so in REM sleep. The mean reliability of the envelope EEG spectrum (pooled across channels, frequency bands and envelope frequencies) was 0.886 (even-odd, SD=0.085) and 0.819 (split-half, SD=0.14) for NREM and 0.513 (even-odd, SD=0.157) and 0.684 (split-half, SD=0.141) for REM. No clear trend was seen by envelope frequency (Figure 4). The reliability of the mid-frequency EEG bands (alpha and low sigma) was the highest, falling off towards both low and high frequencies (Supplementary figure S2), while no clear trend was seen by scalp channel (Supplementary figure S3).

**Figure 4.**
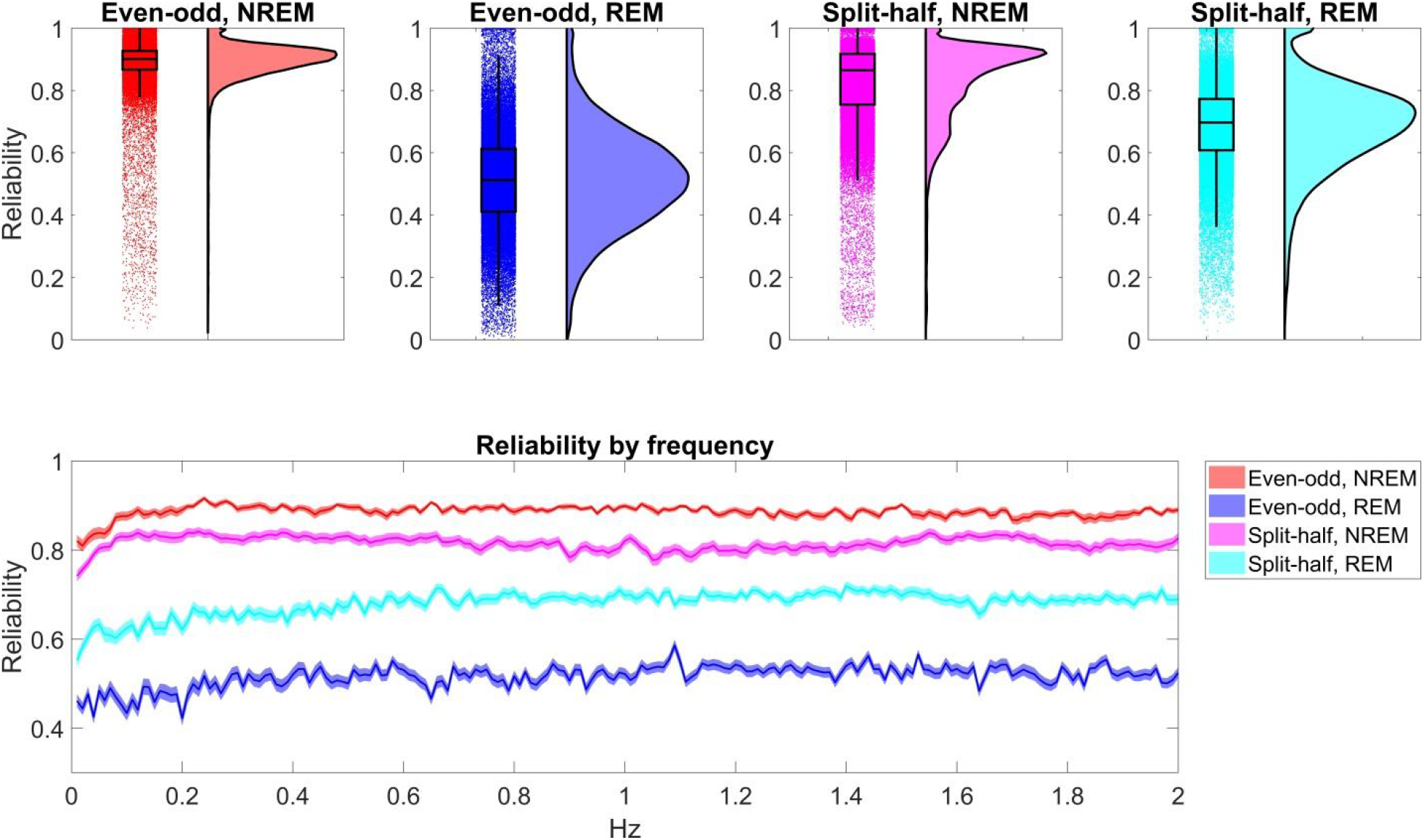
The reliability of the sleep EEG envelope spectrum. Upper panels are raincloud plots (Allen et al., 2019) by vigilance state and reliability type, showing raw data overlain with box plots on the left side and kernel density curves on the right side. Data from all frequency bands, envelope frequency bins and scalp channels are pooled. The lower panel illustrates reliability by envelope frequency bin. Data from all frequency bands and scalp channels are pooled, shading indicates 95% confidence intervals of the mean.

### The envelope spectrum reflects age and sex, but not general cognitive ability

The envelope spectrum of the sleep EEG was significantly associated with demographic variables, but not with general cognitive ability (Figure 5, Figure 6). This association was the strongest between the NREM PSD and age. Older age was generally associated with a loss of low-frequency oscillations in the power of NREM EEG frequencies, often with an increase of higher-frequency oscillations. Specifically, a reduced oscillation of low delta power at ∼0.5 and ∼1.6-1.8 Hz, but increased oscillation at 1–1.5 Hz; a reduced ∼0.25 Hz oscillation of theta, alpha, sigma and beta power with an increased 0.5–1 Hz oscillation of theta and sigma power was seen. In REM sleep, a general tendency for increased low- and high-frequency power oscillations and a corresponding decrease at ∼0.5–1 Hz was seen, but this only reached statistical significance in the high delta, alpha and low sigma frequency bands.

**Figure 5.**
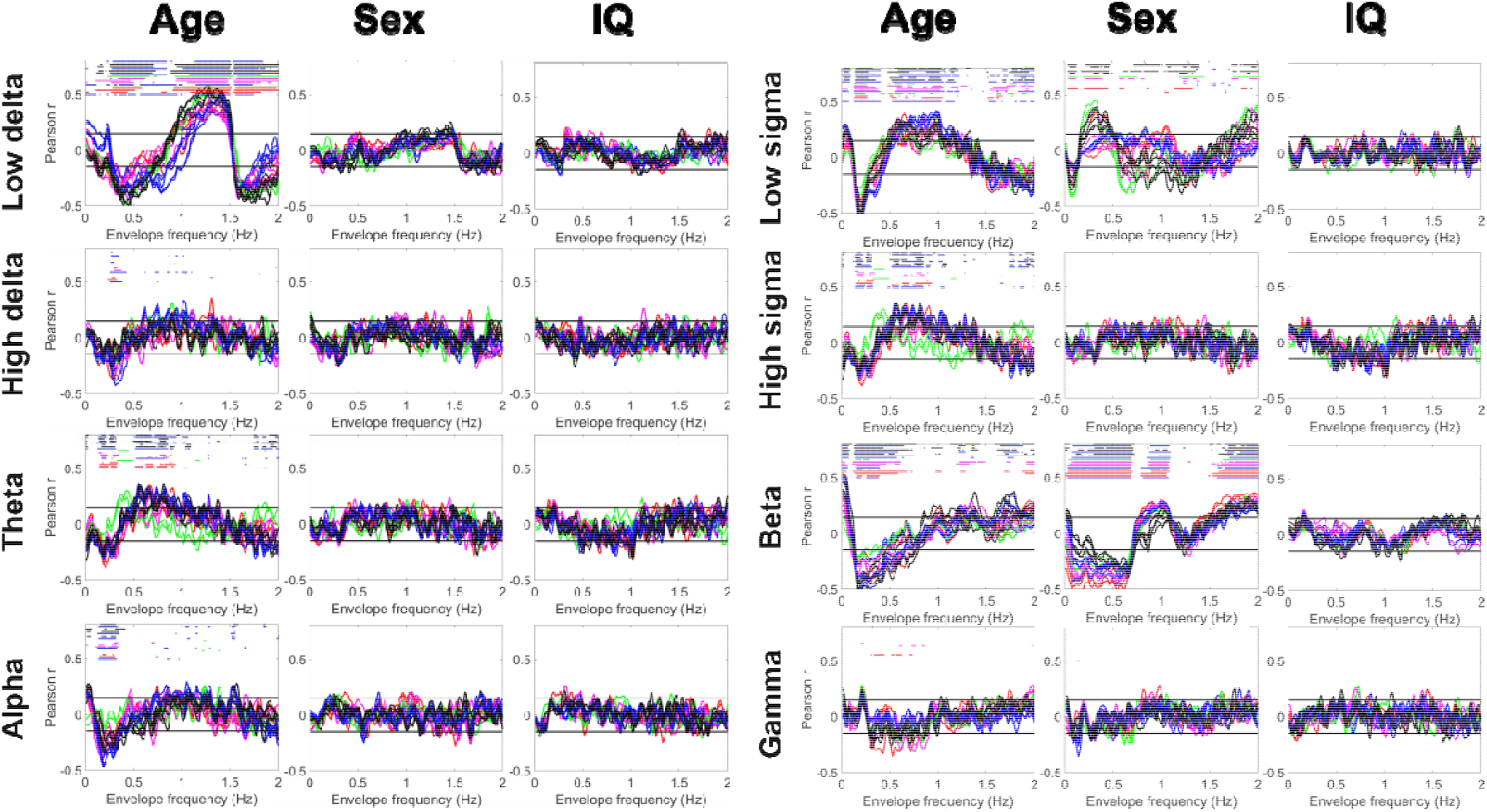
Correlations between the NREM envelope spectrum and age, sex and general cognitive ability (IQ). Colored lines represent correlation coefficients by scalp channel. Color codes indicate scalp region, with individual channels from the same region shown with the same color. Black horizontal lines show the threshold of conventional (p=0.05) significance). Colored dots (with color coding identical to lines) above the lines indicate a statistically significant correlation after FDR correction on the corresponding channel.

**Figure 6.**
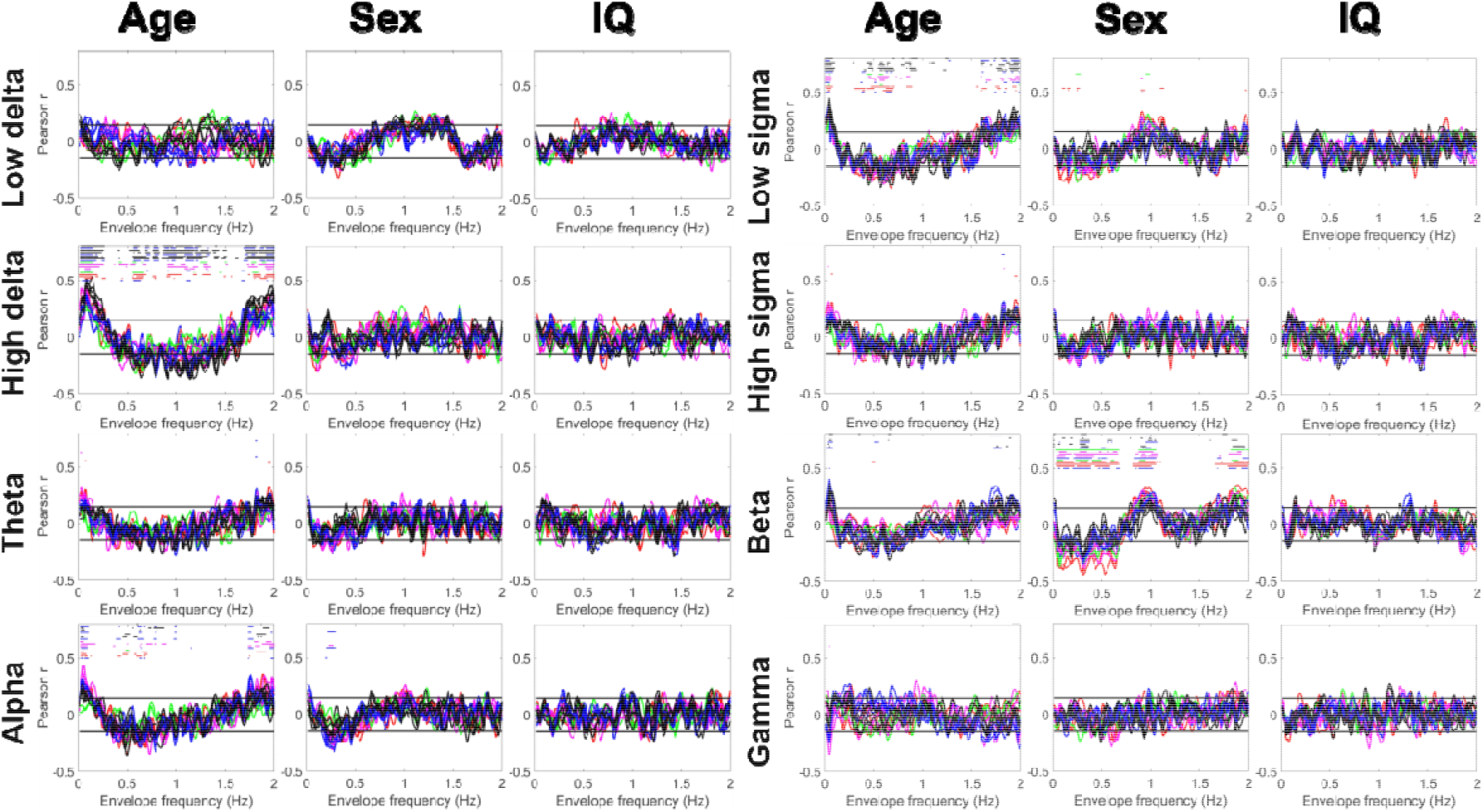
Correlations between the REM envelope spectrum and age, sex and IQ. Colored lines represent correlation coefficients by scalp channel. Color codes indicate scalp region, with individual channels from the same region shown with the same color. Black horizontal lines show the threshold of conventional (p=0.05) significance). Colored dots (with color coding identical to lines) above the lines indicate a statistically significant correlation after FDR correction on the corresponding channel.

Male sex was associated with a lower amplitude of ∼0.05–0.1 and ∼0.5–1.5 Hz NREM low sigma power oscillations, but a higher amplitude of power oscillations of the same frequency band at ∼0.25–0.5 Hz and >1.75 Hz. Male sex was also associated with a lower amplitude of <0.75 Hz and >1.75 Hz, but a higher amplitude of ∼1 Hz beta power oscillation irrespective of sleep state.

General cognitive ability was not significantly associated with the envelope spectrum of either NREM or REM sleep EEG.

### Multivariate models

The relationship between human phenotypes and single biological markers, such as single genetic polymorphisms or individual features of brain morphology is usually modest. However, multivariate models using a large number of such biological markers as independent variables are able to capture the additive, independent contribution of each single marker to reach a much more substantial correlation between the totality of biological markers and phenotypes (Lessov-Schlaggar et al., 2016). Therefore, beyond demonstrating correlations between single spectral features of the sleep EEG envelope and age, sex and intelligence, we set out to investigate the relationship between these features using multivariate models. Because of the modest sample size (N=176) for the very large number of possible features (200 PSD values from 8 bands on 18 channels, separately from NREM and REM sleep), we used elastic net regression, a learning algorithm for training (N=154), with an independent validation sample (N=22), we only used the first 100 PSD bins as these exhibited the largest bivariate correlations with phenotypes, and we ran models separately by channel.

The total predictive validity of the envelope spectrum towards each phenotype was expressed as the correlation between predicted and actual values in the validation sample (predictive accuracy) (Figure 7). Using NREM sleep, age could be predicted with reasonable accuracy (r_mean_=0.616, r_SD_=0.151, the prediction accuracy for sex was lower but still substantial (r_mean_=0.447, r_SD_=0.138), but the correlation between predicted and actual IQ was low (r_mean_=0.151, r_SD_=0.178). Using the REM sleep envelope, age could be predicted with moderate accuracy (r_mean_=0.502, r_SD_=0.156), but this was not the case for sex (r_mean_=-0.019, r_SD_=0.224) and IQ (r_mean_=0.092, r_SD_=0.112). (All means and SDs are across channels.) In case of IQ, elastic net models frequently failed to converge due to the low correlation between PSD values and this phenotype.

**Figure 7.**
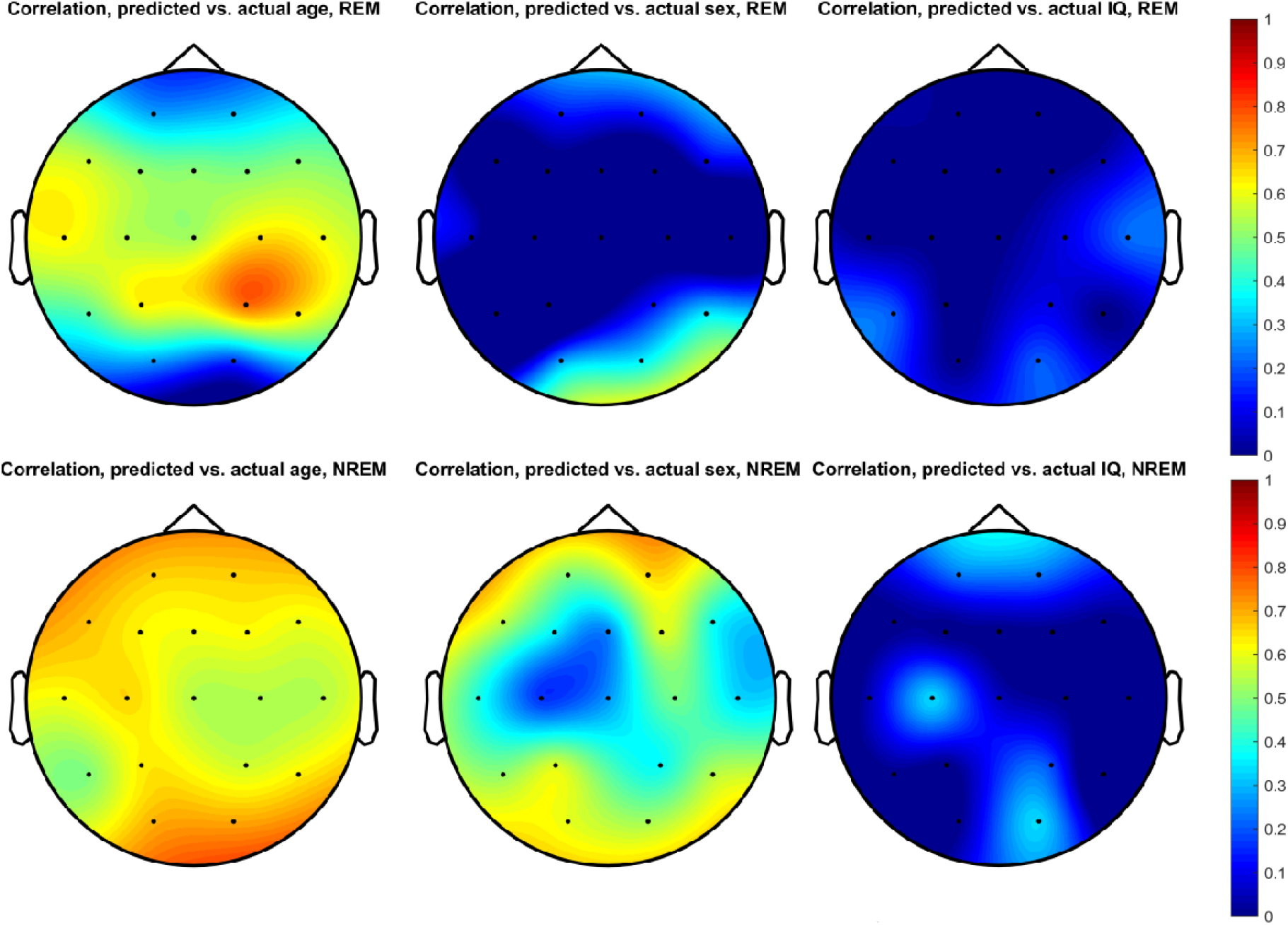
The performance of elastic net regression models predicting age, sex and IQ from the envelope spectrum. Topographic plots illustrate the correlation between predicted and actual phenotypes in the validation sample. (Elastic net regression models were run separately for each channel). The correlation for channels on which the elastic net model did not converge is set to 0 and not counted towards the average performance described in the text.

### Relationship to respiratory rhythms

Low-frequency fluctuations in the EEG signal could theoretically be affected or contaminated by lthe respiratory cycle, which also occurs with sub-second periods. In order to investigate to what extent this occurs, we used the recordings of 29 participants containing a pair of respiration channels to estimate to what extent respiratory activity is correlated with the course of EEG envelopes. We estimated the 1) magnitude-squared coherence between each EEG band envelope and respiratory activity 2) the modulation index between respiratory activity and EEG band envelopes. We performed these calculations with 100 second windows of artifact-free data with 50% overlaps between windows and compared statistics to those calculated from 1000 random surrogates to estimate statistical significance in each participant. Like in other analyses, we transformed the resulting empirical p-values to z-values, averaged them across participants and transformed them back to p-values before application of the Benjamini-Hochberg correction of false discovery rate.

In line with a previous study (Achermann and Borbély, 1997) we found no coupling between EEG envelopes and respiratory activity. Neither coherence between respiratory activity and EEG envelopes nor their modulation index was ever significantly higher than in surrogates, illustrated by an almost perfectly circular phase histograms of EEG envelope amplitudes as a function of respiration phase (Supplementary figure S4-S5). Thus, our findings confirm that EEG amplitude fluctuations occur largely independently from low-frequency respiratory rhythms, and thus the EEG envelope is not a respiratory artifact.

## Discussion

In our study, we aimed to describe the sleep EEG envelope in detail and compare its characteristics to the ordinary sleep EEG spectrum to assess its viability as a biomarker. Overall, our study demonstrates that the sleep EEG envelope shares many of the properties of the ordinary sleep EEG: it reflects neuronal population firing, it has characteristic oscillation frequencies, it is highly individually stable and varies between individuals; and it is associated with demographic characteristics.

It has been shown in previous human invasive EEG studies that sleep oscillations recorded either from the cortex or from the scalp closely reflect rhythmic ensemble firing neuron populations. For instance, slow waves (Csercsa et al., 2010; Nir et al., 2011), sleep spindles (Ujma et al., 2019b) and the wakeful alpha rhythm (Halgren et al., 2019) as field potentials are all associated with waxing and waning patterns in local neuronal firing. We observed a similar pattern for the envelope as well. MUA was significantly suppressed when low-frequency activity was high: specifically, the lowest MUA was observed during the maximum of low delta activity and slightly after the maximum of high delta and theta activity. Curiously, the opposite pattern (increased MUA during periods of reduced low-frequency oscillations) was less typical. This phenomenon may reflect the rhythmic suppression of neuronal firing during slow oscillations (Cash et al., 2009; Csercsa et al., 2010; Nir et al., 2011), which contain ensembles of low frequencies up to the alpha range (Borbely et al., 1981). Although such neuronal down-states are generally followed by up-states containing high frequency rhythms (Clemens et al., 2007; Muehlroth and Werkle-Bergner, 2020), the fact that down-states are generally more prominent (Cash et al., 2009; Nir et al., 2011) may specifically result in an association between the presence of low-frequency activity in the ECoG and reductions in neuronal firing in nearby cortex. For high frequencies (low sigma through gamma, with alpha being an intermediate range), an opposite pattern was seen: MUA was maximal when oscillations in these frequencies were gaining in power, possibly reflecting the role of cortical neuronal assemblies in recruiting these oscillations.

Next, we used scalp EEGs for healthy volunteers to establish further properties of the sleep EEG envelope spectrum. We found that, similarly to the ordinary spectrum, the envelope spectrum was also characterized by higher powers at lower frequencies. In line with previous reports on slow wave and sleep spindle periodicity (Kubicki et al., 1986; Achermann and Borbély, 1997) we found two characteristic peaks: one at ∼0.05 Hz (20 second period, most prominent for slow rhythms), and another at ∼0.25 Hz (4 second period, most prominent for faster rhythms). These frequency peaks were less prominent in REM sleep than in NREM.

Previous reports have established that the sleep EEG spectrum is fingerprint-like with a high intra-individual stability (Tan et al., 2000; Finelli et al., 2001; Tan et al., 2001; De Gennaro et al., 2005; Reynolds et al., 2018), which is the result of genetic regulating factors (Ambrosius et al., 2008; De Gennaro et al., 2008; Landolt, 2011; Adamczyk et al., 2015). Although our ability to fully replicate this finding in the envelope spectrum was limited by the absence of multiple recordings and genetically informative data, we could establish that when comparing spectra from the same individual across the two halves of the night or across even and odd numbered sampling windows, reliability was very high for NREM (>0.8) and reasonably high for REM (>0.5), with remarkably similar reliability values across all but the lowest frequencies.

The reliability of the sleep EEG envelope spectrum renders it a potential marker of stable individual differences, such as demographic variables, psychological traits or pathological conditions. In a quantitative test of this hypothesis, we found that higher age was associated with reductions in the 0.25 Hz rhythmicity of high delta through beta rhythms. A relative increase in the ∼1 Hz rhythm of sigma-frequency oscillations, an additional increase in the very low frequency rhythms of low sigma and beta oscillations, as well as a relative reduction of low-frequency and a relative increase of high-frequency low delta oscillations was also seen. These results – together with findings from invasive EEG recordings – can be interpreted as a systematic loss of the medium-scale temporal organization of rhythmic neuronal firing as a function of ageing. Notably, envelope spectra calculated from NREM were much more associated with age than REM spectra, highlighting the functional importance of this vigilance state for ageing-related phenomena (Mander et al., 2013; Mander et al., 2017; Sun et al., 2019; Ujma et al., 2019a).

Sex was associated with a single EEG envelope feature: low-frequency rhythmicity of the NREM beta rhythm was reduced in males, while high-frequency rhythmicity was higher. The significance of the ∼1 Hz rhythm suggests that beta rhythms show stronger coupling to slow waves in males, however, the functional importance of this finding is currently unknown.

Although intelligence was found to be associated with multiple sleep EEG spectral features (Ujma et al., 2017; Ujma, 2018), we found no evidence that it is also associated with the rhythmicity of sleep EEG oscillations.

We used a learning algorithm to perform multivariate predictions of age, sex and intelligence based on the sleep EEG envelope spectra. As expected based on the reliability of spectra, much better predictions could be made based on NREM than REM spectra. Age could be predicted with reasonable accuracy from NREM sleep (r∼0.6), although much more accurate predictors were previously constructed based on overall features of the sleep EEG (Sun et al., 2019) or the shape of NREM slow waves (Ujma et al., 2019a). Sex could be predicted from the NREM envelope spectrum with lower but still substantial accuracy (r∼0.45), although the predictive power of the REM spectrum was much lower. Sex prediction based on the envelope spectrum underperforms relative to other predictors based e.g. on brain imaging (Sepehrband et al., 2018; Anderson et al., 2019; Dhamala et al., 2020). However, we did not expect the envelope spectrum to be a particularly sexually dimorphic characteristic. The non-significant zero-order correlations between the envelope spectrum and intelligence could not be improved with the use of elastic net regression: models failed to converge on most electrodes and even with this method we found no association between the envelope spectral and intelligence.

What biological process do amplitude fluctuations in the sleep EEG reflect? In Table 1 we provide a non-exhaustive list of known biological oscillations with periods at most on the minute scale. From this list, we had data about two prominent oscillations: the cardiac and the respiratory rhythm. Both oscillations could theoretically drive low-frequency EEG rhythms either through physiological mechanisms (for example, because neuronal firing depends on the availability of oxygenated blood and this is reflected in EEG rhythms) or through electrical artifacts detected by the EEG. However, based on non-significant magnitude-squared coherence and phase-amplitude coupling the respiratory rhythm appears to play a role in low-frequency EEG amplitude oscillations, and the cardiac rhythm is too fast to strongly influence all but the fastest envelope rhythms. Because our recordings did not contain data about other oscillating biological processes, we can only speculate about their role. With their characteristic 20-second periods, gastric rhythms (Wolpert et al., 2020) oscillate at a frequency strongly overlapping with characteristic envelope frequencies, rendering EEG envelope oscillations a promising potential marker in the study of brain-viscera interactions. Other known biological rhythms are not strong candidates to be the driver of or to be coupled with EEG amplitude oscillations due to differences in their characteristic frequencies. In sum, the precise biological mechanism creating periodic fluctuations in the amplitude of EEG rhythms remains unknown and its discovery is a major task of further studies into this phenomenon.

**Table 1.** Biological processes with low-frequency oscillations. The list of oscillations and data on their characteristics are from (Goldbeter, 1997) unless otherwise indicated.

| Oscillation (reference) | Period | Frequency |
| --- | --- | --- |
| Main cardiac rhythm | 1 second | 1 Hz |
| Respiratory rhythm | 4 seconds | 0.25 Hz |
| Calcium oscillations | 1 second – several minutes | <0.016-1 Hz |
| Resting state alpha power<br>(Omata et al., 2013) | 6–100 seconds | 0.01-0.17 Hz |
| Gastric rhythms (Wolpert et al., 2020) | ~20 seconds | ~0.05 Hz |
| Hormonal rhythms | At least several minutes | <0.016 Hz |
| Cell cycle | 10 minutes – 1 day | <0.001 Hz |

Our work has a number of limitations. First, using a single IME per patient we were only able to record neuronal firing from a very limited cortical area. EEG recorded on the adjacent cortical surface is likely the summation of neuronal activity in a more extended area, consequently, the correlation between MUA and the envelope was not particularly strong. Second, we had only a single night of measurement from healthy individuals, resulting in a within-night, rather than a more optimal across-night estimation of envelope reliability.

In sum, our study revealed that the periodicity of amplitude fluctuations in the sleep EEG, reflected by the envelope, is a promising human biomarker. In an invasive study, it was found to be associated with fluctuations in neuronal firing. In a study of healthy volunteers, it was found to be a highly reliable individual marker, somewhat sexually dimorphic and especially strongly associated with ageing. While we showed that envelope fluctuations reflect fluctuations in neuronal firing, why these fluctuations take place (and why they change with ageing) requires further study.

## Data availability

Supplementary data and MATLAB codes are available on Zenodo at: 10.5281/zenodo.5595341

Raw EEG data is available upon reasonable request to the corresponding author.

## Acknowledgements

This article is based upon work supported by the National Research, Development and Innovation Office of Hungary (grants NKFI_FK_128100, K_128117, NKFIH-1157-8/2019-DT and 2017-1.2.1-NKP-2017-00002), the Higher Education Institutional Excellence Program of the Ministry of Human Capacities in Hungary within the framework of the Neurology thematic program of the Semmelweis University, a Vidi grant from the Netherlands Organisation for Scientific Research (NWO), and COST Action CA18106 supported by COST (European Cooperation in Science and Technology).

