## Supplementary figure S1 for "The sleep EEG envelope: a novel, neuronal firing-based human biomarker"

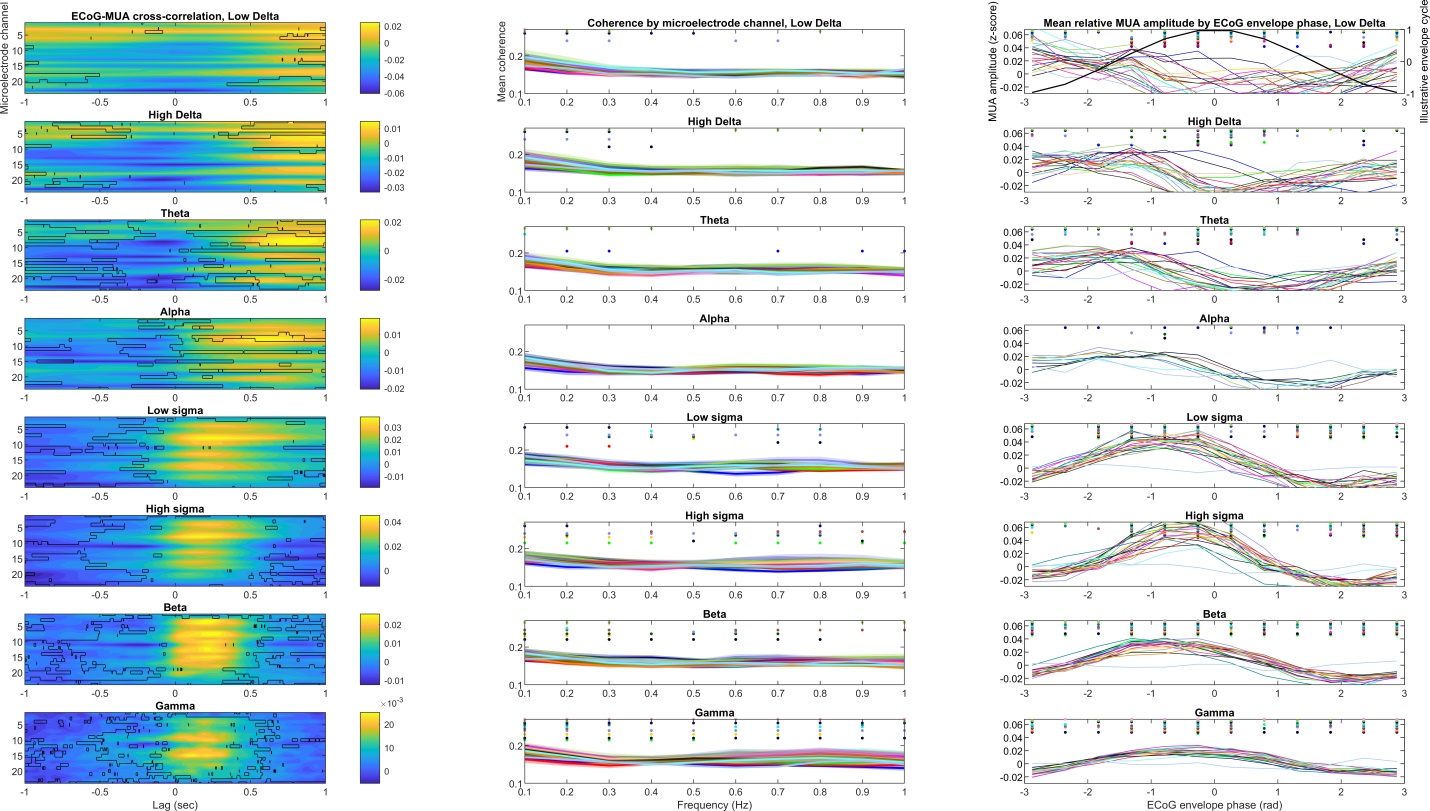

Supplementary figure S1. Coupling between EEG envelope in the cortical surface and MUA within the adjacent cortex in NREM sleep, with the exclusion of three patients with potentially artefactual data. **Panel A**: cross-correlation of the two signals. The horizontal axis indicates time lags, the vertical axis indicates IME channel (N=23, deeper channels shown at the bottom), while the color axis indicates correlation coefficients. Black outlines show statistically significant results after FDR correction. **Panel B**: magnitude-squared coherence between the two signals. Overlain lines represent individual IME channels. Because of the large number of channels and no substantial between channel differences, no particular pattern in color coding was used. Dots indicate statistical significance after FDR correction on the corresponding channel. Deeper channels are shown at the top. **Panel C:** Mean MUA amplitude (in within-segment z-scores) by ECoG envelope phase bins. Dots indicate statistical significance after FDR correction on the corresponding channel. A sinusoid is overlain in the low delta subplot for illustration. On panel B and C, for better visibility only IME channels are shown where at least one data point reached significance.

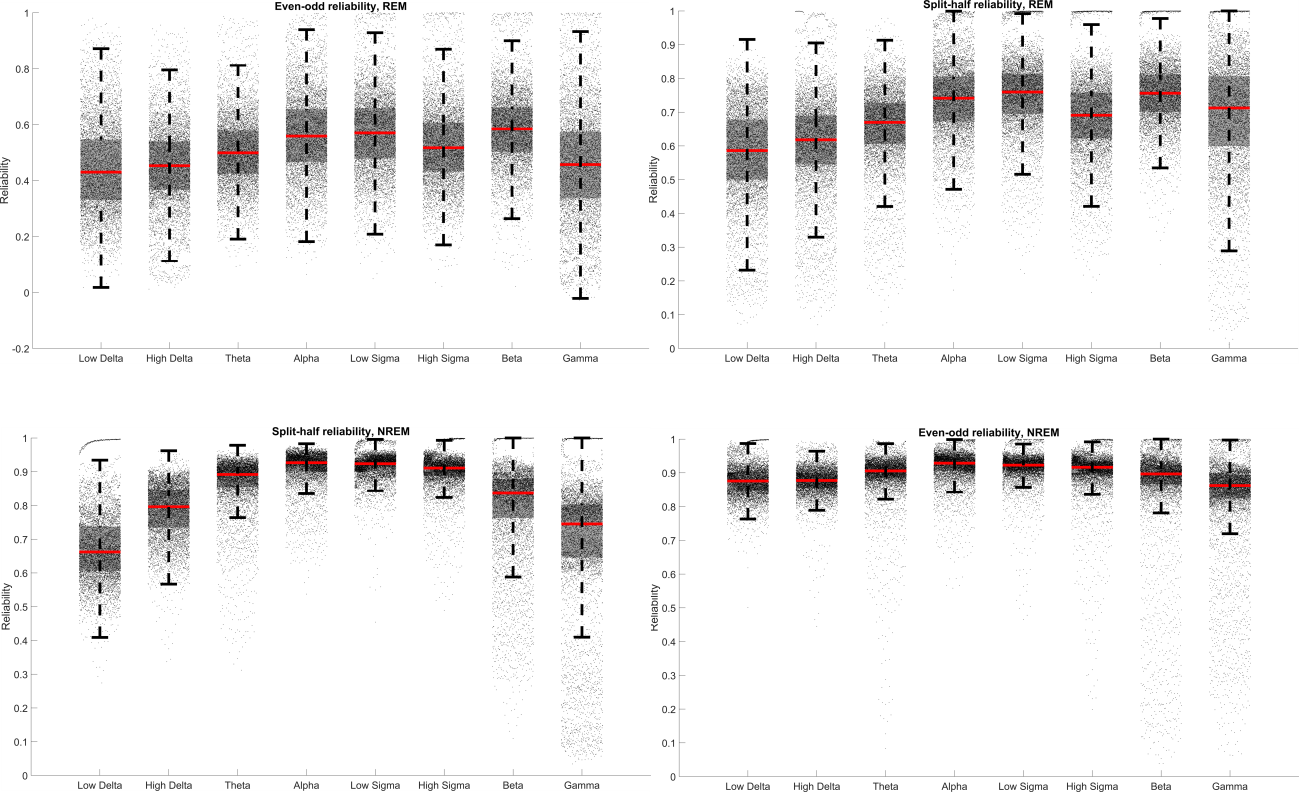

Supplementary figure S2. Reliability estimates by frequency band. Data is pooled across envelope frequency bins and scalp channels and shown as a boxplot. Red lines indicate medians, gray boxes enclose the 2^nd^ and 3^rd^ quartiles, while whiskers show 1.5 interquartile ranges.

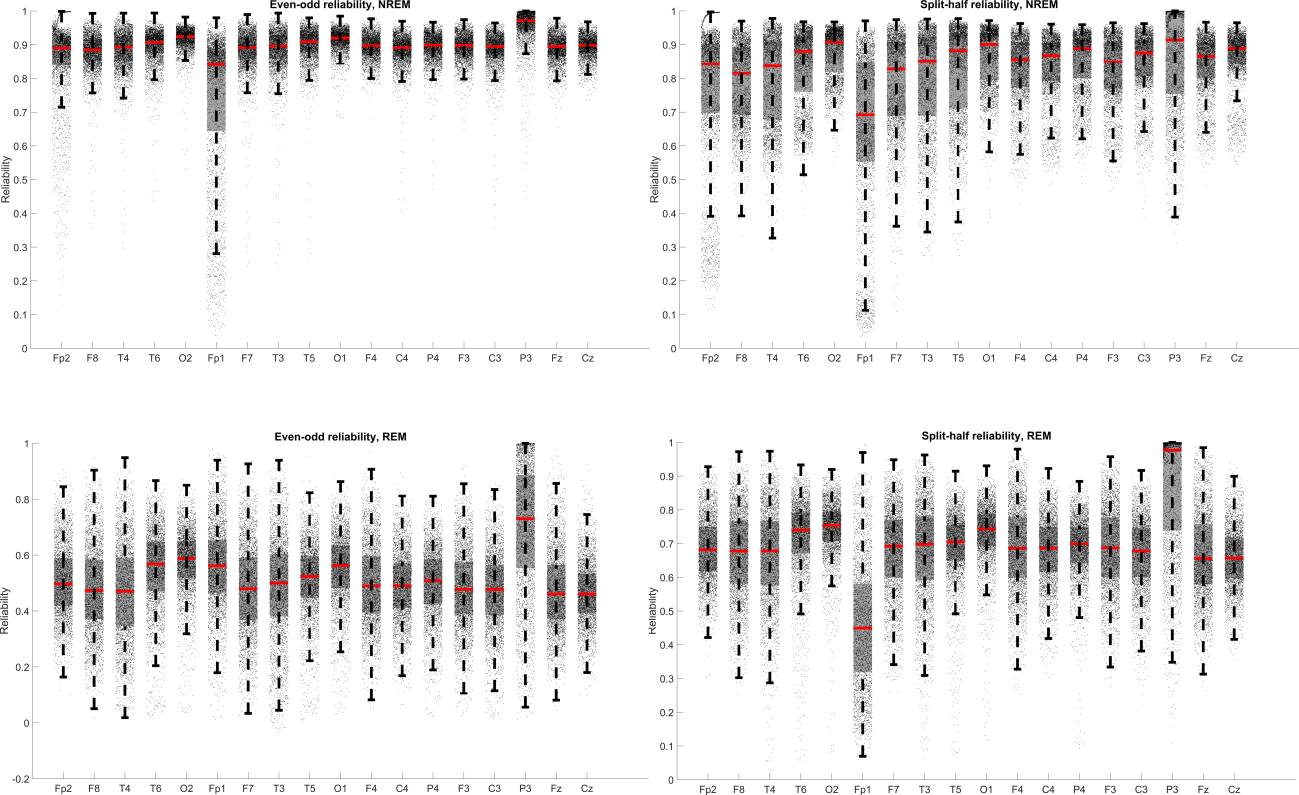

Supplementary figure S3. Reliability estimates by scalp channel. Data is pooled across envelope frequency bins and frequency bands and shown as a boxplot. Red lines indicate medians, gray boxes enclose the 2^nd^ and 3^rd^ quartiles, while whiskers show 1.5 interquartile ranges.

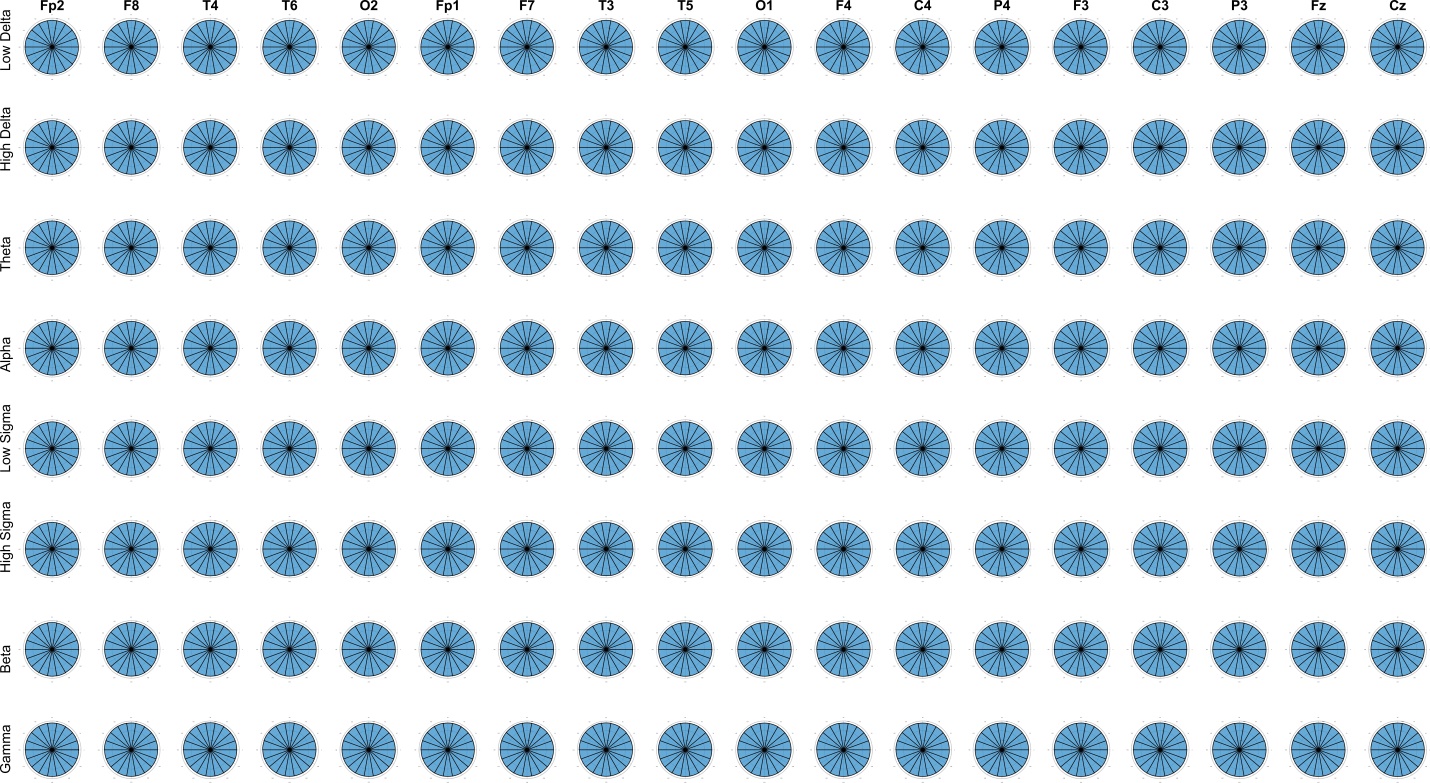

Supplementary figure S4. Circular histograms showing the distribution of EEG power across 18 equally spaced phase bins of the respiratory cycle in NREM. For better visibility we removed all captions from the histograms. Note that the circular histograms are almost perfectly round for each channel and each frequency band, indicating no phase-frequency coupling.

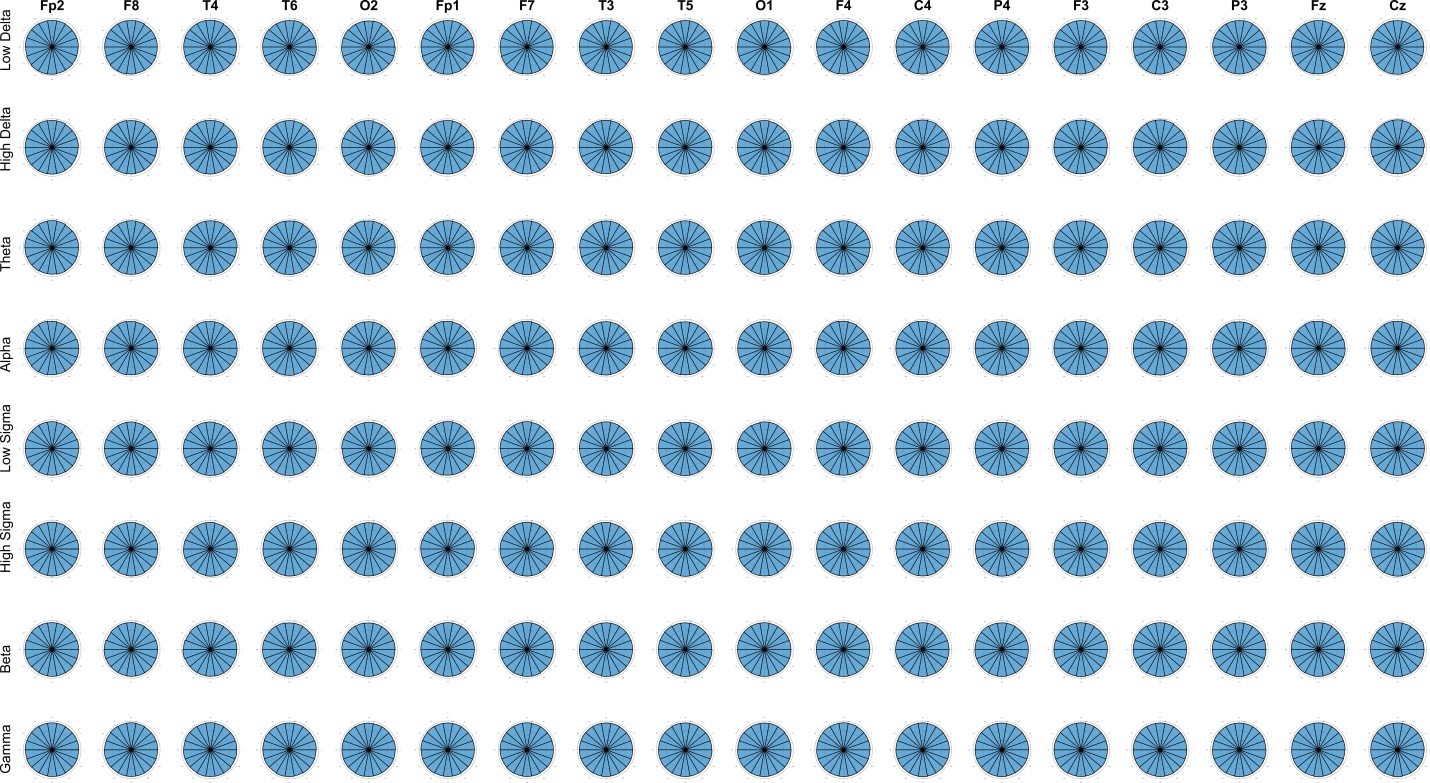

Supplementary figure S5. Circular histograms showing the distribution of EEG power across 18 equally spaced phase bins of the respiratory cycle in REM. For better visibility we removed all captions from the histograms. Note that the circular histograms are almost perfectly round for each channel and each frequency band, indicating no phase-frequency coupling.
